# Sympathetic-to-microglia readouts in human brain single-nucleus data: sparse adrenergic receptor detection and age-associated homeostatic-to-inflammatory shifts in hippocampal microglia

**DOI:** 10.64898/2026.11.14.765300

**Authors:** Min Liang, Kai Zeng

## Abstract

Stellate ganglion block (SGB) improves sleep in clinical populations, yet the mechanisms linking sympathetic activity to human microglial states remain untested. A prerequisite is to establish what sympathetic signaling readouts exist in human microglia and whether state composition changes in a disease-relevant physiological dimension. Using two public human single-nucleus datasets (the ROSMAP prefrontal cortex resource: 443 donors, 174,420 immune nuclei, 152,459 microglial nuclei, 12 published microglial states; and a hippocampal multiome cohort spanning adult lifespan: 40 donors, ages 20–100, 15,225 microglial nuclei), we quantified (i) adrenergic receptor and inflammatory/complement axis gene expression across microglial states, and (ii) donor-level trajectories of four microglial state signatures across age. Adrenergic receptors were barely detected: among nine receptor genes, only *ADRB2* was detectable in 3.8–11.4% of nuclei, and no adrenergic receptor was state-specific. In contrast, inflammatory genes (nuclear factor-κB and interleukin modules) were strongly concentrated in one pro-inflammatory state (e.g., *NFKB1* mean expression 1.47 vs 0.35–0.72 in other states; *IL1B* in 32.6% of nuclei vs 2–16% elsewhere), complement genes in a complement-enriched state, and lipid-metabolism/disease-associated genes in a distinct state. Across the adult lifespan, hippocampal microglia showed a donor-level decline of the homeostatic signature (mean score 0.70 to 0.38 across age quartiles; Spearman ρ = -0.55, P = 2.6 × 10^-4^), with concordant rises of pro-inflammatory (ρ = 0.37, P = 0.017) and disease-associated/lipid-metabolism (ρ = 0.32, P = 0.045) signatures; an interferon signature showed no age trend. These results (a) indicate that human microglia lack a robust direct adrenergic receptor substrate at single-nucleus resolution, implying that sympathetic influence on microglia, if causal, is likely indirect or low-frequency, and (b) establish a donor-level, age-associated baseline shift toward inflammatory/trimming states in the human hippocampus. Together they define candidate readouts and the baseline against which chronic-sleep-disorder and neuromodulation-intervention studies (e.g., SGB) should be designed.

## 1. Introduction

Stellate ganglion block (SGB), a cervical sympathetic block used clinically for sleep disturbance, has documented clinical effects on subjective and objective sleep measures in controlled trials, but its central nervous mechanisms are unknown. A plausible mechanistic chain runs from sympathetic tone to hippocampal microglial states, and from microglial states to sleep architecture and cognition. Testing this chain requires, first, an answer to a prior question that has not been addressed systematically in human data: **what is the receptor-level substrate for sympathetic signaling in human microglia, and in which microglial states do inflammatory and trimming modules reside that sympathetic activity could, directly or indirectly, modulate?**

Two public human single-nucleus resources allow a first, donor-level answer:

1. **ROSMAP prefrontal cortex microglia states**: 12 microglial states defined in 152,459 nuclei from 443 community-dwelling aging donors, with published state-specific marker tables and a raw immune-nucleus count matrix available for independent verification.
2. **A hippocampal multiome cohort spanning the adult lifespan**: 40 donors aged 20–100, enabling donor-level tests of whether microglial state signatures shift across age, i.e., whether a homeostatic-to-inflammatory baseline shift exists in the structure in which sleep-disorder mechanisms are typically interrogated.

We asked three questions: (i) Are adrenergic receptors expressed, and are they state-specific, in human microglia? (ii) Where in the state space do inflammatory (NF-κB/interleukin), complement, lipid-metabolism (disease-associated), and interferon modules reside? (iii) Do donor-level state-signature trajectories shift across adult age in human hippocampus? We report the analyses as a methods-transparent baseline reference for studies of neuromodulation (including SGB) and chronic sleep disorders in animal models, and pre-register the interpretation limits inherent to single-nucleus detection of low-abundance transcripts.

## 2. Results

### 2.1 Adrenergic receptors are barely detected in human microglia and are not state-specific

We quantified nine adrenergic receptor genes (*ADRA1A–D, ADRA2A–C, ADRB1–2*; *ADRB3* was not present in the 16,228-gene count-matrix subset) in 152,459 microglial nuclei across the 12 published states (CP10K + log1p normalization).

- **α**_**1**_ **and α**_**2**_ **receptors were effectively undetected**: *ADRA1A/ADRA1D/ADRA2A/ADRA2B* mean normalized expression ≈ 0.00, with < 0.3% of nuclei showing detectable expression in any state.
- ***ADRB2* was the only receptor with appreciable detection**: mean 0.06–0.16 across states, expressed in 3.8–11.4% of nuclei (highest in the homeostatic state MG0, 11.4%; lowest in MG1, 3.8%). *ADRB1* and *ADRA2C* were detected in only 1–2% of nuclei.
- **No adrenergic receptor appeared in the published state-specific marker table** (12 states × 2,228 genes): receptor expression is, at best, a global low-level feature rather than a state-defining one.

These results caution against assuming a robust direct receptor substrate for norepinephrine in human microglia. Given known single-nucleus under-detection of low-abundance transcripts, and the possibility of signal relay through vascular, meningeal, or other glial elements, the results do not exclude sympathetic influence on microglia; they define its plausible magnitude and route as **indirect or low-frequency**, a hypothesis that in-vivo perturbation must test.

### 2.2 Inflammatory, complement, and disease-associated modules occupy distinct states

Mapping 38 axis genes (adrenergic, inflammatory, complement/trimming, interferon) onto the 12-state space showed strong, reproducible state concentration of inflammatory and trimming programs:

- **Pro-inflammatory state (MG10)**: *NFKB1* mean 1.47 (vs 0.35–0.72 in all other states), *IL1B* 0.79 with 32.6% of nuclei positive (vs 2–16% elsewhere), *NFKBIA* 0.78, *TLR2* 1.88. In state-specific log2FC terms, *NFKB1* = 1.60, *IL1B* = 2.22, *NFKBIA* = 1.04.
- **Complement-enriched state (MG3)**: *C1QA/C1QB/C1QC* 0.86–0.89, *APOE* 0.81.
- **Lipid-metabolism / disease-associated state (MG4)**: *TREM2, ITGAX* (0.63), *APOE*, with *SPP1* concentrated in MG7 (1.10).
- **Homeostatic states (MG0/MG1)**: *P2RY12* (0.69) and *CX3CR1* (0.57) anchored to MG0; the loss of these same receptors is described as a priming signature in chronic-stress literature.
- **Interferon state (MG11)** and DNA-damage state (MG12) each formed their own compartments.

Notably, *NLRP3* itself was weakly expressed at human baseline (mean 0.18–0.30; 8–21% of nuclei), and *TLR4, TNF*, and *IL6* were absent from the count-matrix gene subset — a reminder that inflammasome-centric narratives established in rodents or pathological tissue should not be projected onto the human baseline without verification.

### 2.3 Hippocampal microglia shift from homeostatic toward inflammatory/lipid-metabolism signatures across the adult lifespan

In the hippocampal multiome cohort (40 donors, four age strata of 10 each; 15,225 microglial nuclei), we scored four state signatures per nucleus (mean of normalized expression over signature genes; CP10K + log1p) and aggregated to donor level (primary inference unit; pooled cell-level statistics are reported only as sensitivity checks, since they inflate effective n).

| Signature (representative genes) | ρ vs age (donor-level, n = 40) | P | Mean score 20–39 to 80–100 |
|---|---|---|---|
| Homeostatic (*P2RY12/TMEM119/CX3CR1/SALL1/GPR34/OLFML2B/CXXC5*) | **-0.55** | **2.6 × 10<sup>-4</sup>** | 0.70 to 0.51 to 0.38 to 0.39 |
| Pro-inflammatory (*IL1B/TNF/NFKB1/CCL2-4/TLR2/S100A8/A9/NFKBIA/CD83*) | **+0.37** | **0.017** | 0.17 to 0.18 to 0.21 to 0.26 |
| DAM/lipid-metabolism (*TREM2/APOE/C1QA-C/C1R/C1S/ITGAX/CLEC7A/SPP1/LPL/C3*) | **+0.32** | **0.045** | 0.61 to 0.58 to 0.85 to 0.72 |
| Interferon<br>(*ISG15/IFIT1-3/MX1/OAS1/OASL/IRF7/IFI6*) | -0.23 | 0.148 (ns) | mild decline |

The homeostatic decline was monotonic across the three younger strata and plateaued in the oldest; the pro-inflammatory rise was strictly monotonic; the DAM/lipid-metabolism signature peaked in the 60–79 stratum. Pooled cell-level Spearman correlations were directionally concordant (homeostatic -0.29; pro-inflammatory +0.18; DAM +0.22; all P < 10^-100^) but should be interpreted only as sensitivity analyses given n-inflation.

These data establish, at donor level, that **the human hippocampal microglial state composition shifts away from homeostasis toward inflammatory/lipid-metabolism programs across adult age**, a physiological baseline against which chronic sleep disturbance, sympathetic activity, and neuromodulation effects must be measured.

## 3. Discussion

### Implication 1: the direct-receptor route is weak, and perturbation studies are mandatory

The near-absence of detectable adrenergic receptors in human microglia implies that any sympathetic effect on microglial states, such as the clinically observed benefits of SGB on sleep, is unlikely to be explained by a robust direct norepinephrine-to-microglia receptor substrate. Candidate indirect routes (vascular and meningeal interfaces, other glial elements, systemic cytokine mediators) require in-vivo perturbation designs rather than receptor-centric interpretations of bulk data.

### Implication 2: the module map defines candidate readouts

Because inflammatory (NF-κB/interleukin), complement, and lipid-metabolism programs are concentrated in discrete states, donor-level scores of these modules — rather than binary “activation” — are the natural readouts for intervention studies. The co-location of *P2RY12/CX3CR1* loss with priming phenotypes in the chronic-stress literature links our human baseline map to experimental perturbation literature in animal models.

### Implication 3: an age-anchored baseline exists in human hippocampus

Chronic sleep disorders in middle-aged and older populations would act on, not in place of, this baseline. Intervention studies should therefore report state-signature shifts relative to age-adjusted expectations rather than to a single “healthy” reference.

### Limitations

(i) Single-nucleus RNA-seq under-detects low-abundance transcripts; receptor “non-detection” is not proof of absence, and the count matrix available for independent verification was a 16,228-gene subset (excluding, e.g., *ADRB3, TLR4, TNF, IL6*). (ii) The ROSMAP cohort is disease-enriched (aging with and without dementia); receptor and module patterns may differ in young healthy tissue. (iii) State annotations were taken from the published resource with GO-term-based naming; formal reconciliation with the original figure-level annotations is warranted. (iv) Signature gene lists were defined a priori from the literature and the published marker tables; alternative signature definitions may shift effect sizes. (v) Donor-level inference (n = 40) is conservative relative to pooled cell-level statistics, which we report only as sensitivity checks.

## 4. Methods

### 4.1 Data sources

- **ROSMAP microglia states**: processed data from the published 12-state resource [1] (metadata for 174,420 immune nuclei; 12-state assignments and StateMarkers table; state GO terms; raw immune-nucleus count matrix, 632 MB RDS). The count matrix was parsed independently with a purpose-built positional parser for RDS dgCMatrix objects; reconstruction was verified by cross-checks (non-zero count, dimensions, byte-level tail alignment). Analyses used the 152,459 microglial nuclei subset and the 16,228-gene × 174,420-nucleus count matrix.
- **Hippocampal multiome cohort (GSE278576)**: Seurat object (31,485 genes × 295,033 nuclei; 40 donors in four age strata of 10 each; downloaded, integrity-verified at byte level, and opened with h5py). The cohort was published as Zemke et al., *Science* 2026 [2]; our analyses here use only the RNA layer of the multiome data.

### 4.2 Preprocessing and scoring

Normalization: counts per 10,000 (CP10K) + log1p. State signatures: fixed a-priori gene lists (Section 2.3 table; plus axis gene panels in Section 2.1–2.2); per-nucleus signature score = mean normalized expression over detected signature genes; donor-level score = mean over the donor’s microglial nuclei. Adrenergic receptor quantification: mean normalized expression and percent-detectable per state.

### 4.3 Statistics

Primary inference at donor level: Spearman rank correlation between donor signature score and age (n = 40); no multiple-comparison correction was applied to the four prespecified signature tests (all were prespecified; nominal P reported). Cell-level pooled correlations reported as sensitivity analyses only. Effect sizes reported as mean scores per age stratum.

### 4.4 Code and data availability

All input datasets are public: the ROSMAP microglia-state resource via its published distribution (Synapse syn52293417; GEO GSE227223), and the hippocampal multiome cohort via GEO (GSE278576). Analysis code (RDS dgCMatrix parser; signature scoring with donor-level aggregation; figure scripts) is archived on Zenodo (DOI: 10.5281/zenodo.24081469). Key intermediate results (donor-level signature scores, per-state gene-expression tables) are included in the code deposit.

### 4.5 Figures

- Figure 1 (composite): (A) axis-gene × 12-state heatmap (ROSMAP; state annotations from the published resource); (B) hippocampal microglia state-signature scores vs age (GSE278576; donor-level Spearman, n = 40); (C) adrenergic receptor percent-detectable bar chart.
- Figure 2: donor-level aging-baseline analysis detail (four signatures × age strata; scatter and trend views).

**Figure 1.**
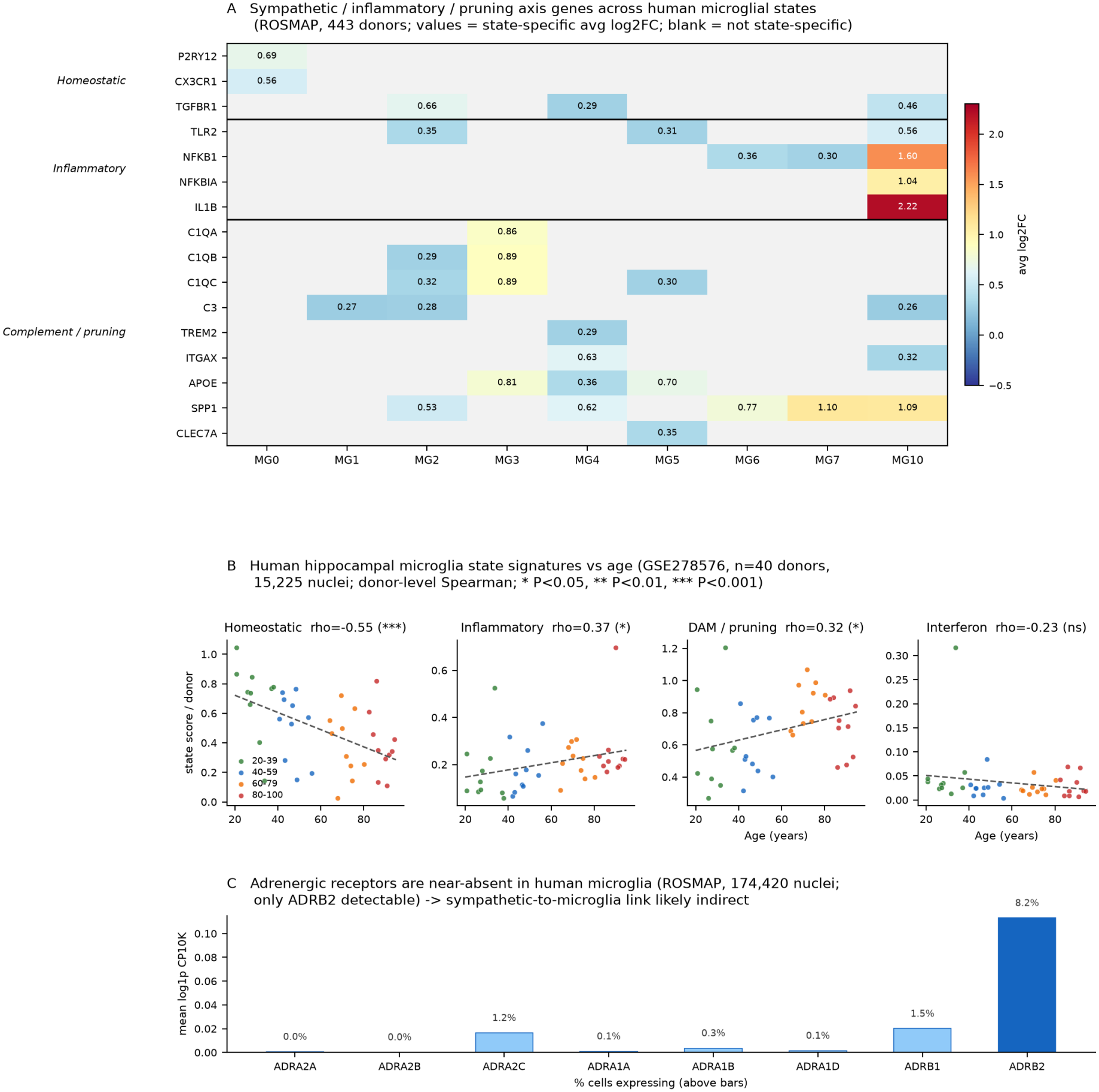
Human microglial sympathetic-to-inflammatory readouts. (A) Axis-gene expression across the 12 published microglial states (ROSMAP, 152,459 microglial nuclei; state annotations from the published resource). (B) Hippocampal microglia state-signature scores vs age (GSE278576, 40 donors; donor-level Spearman; * P<0.05, ** P<0.01, *** P<0.001). (C) Percentage of nuclei with detectable expression of each adrenergic receptor gene, by state.

**Figure 2.**
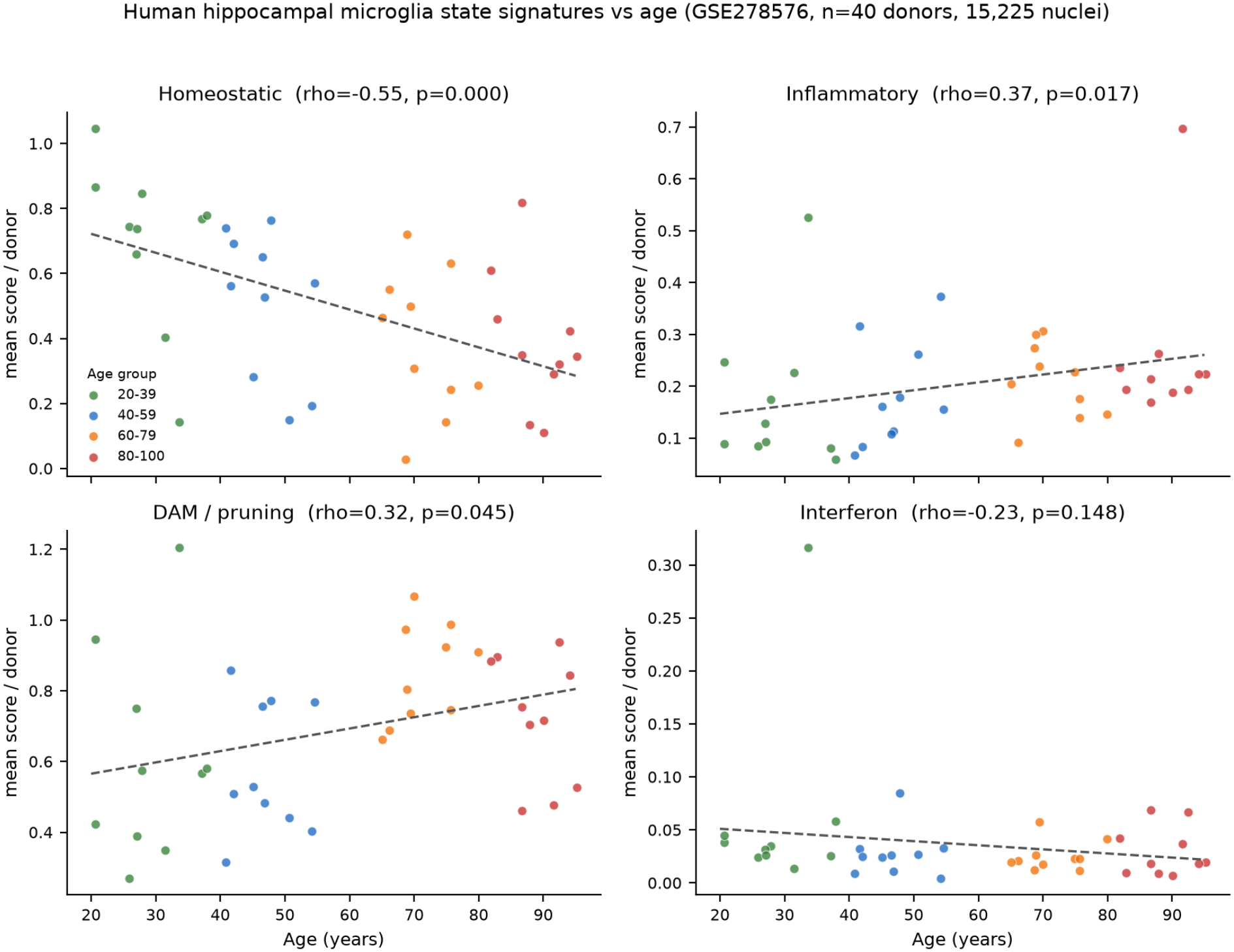
Donor-level aging-baseline analysis of hippocampal microglia state signatures (GSE278576, n = 40 donors, four age strata). Homeostatic signature declines across age strata (0.70 to 0.38), with concordant rises of pro-inflammatory and DAM/lipid-metabolism signatures; interferon signature shows no age trend.

## Acknowledgements

We thank the ROSMAP study participants and the authors of the hippocampal multiome cohort for making data publicly available.

## Author contributions

M.L.: conceptualization, data acquisition, all analyses, code, figures, writing (original draft). K.Z.: supervision, resources, writing (review & editing), endorsement.

